# Annotating and prioritizing human non-coding variants with RegulomeDB

**DOI:** 10.1101/2022.10.18.512627

**Authors:** Shengcheng Dong, Nanxiang Zhao, Emma Spragins, Meenakshi S. Kagda, Mingjie Li, Pedro Assis, Otto Jolanki, Yunhai Luo, J Michael Cherry, Alan P Boyle, Benjamin C Hitz

**Author notes:** These authors contributed equally (Shengcheng Dong, Nanxiang Zhao). Co-Correspondence.

## Abstract

Nearly 90% of the disease risk-associated variants identified from genome-wide association studies (GWAS) are in non-coding regions of the genome. The annotations obtained from analyzing functional genomics assays can provide additional information to pinpoint causal variants, which are often not the lead variants identified from association studies. However, the lack of available annotation tools limits the use of such data.

To address the challenge, we have previously built the RegulomeDB database for prioritizing and annotating variants in non-coding regions^1^, which has been a highly utilized resource for the research community (Supplementary Fig. 1). RegulomeDB annotates a variant by intersecting its position with genomic intervals identified from functional genomic assays and computational approaches. It also incorporates those hits of a variant into a heuristic ranking score, representing its potential to be functional in regulatory elements.

Here we present a newer version of the RegulomeDB web server, RegulomeDB v2.1 (http://regulomedb.org). We improve and boost annotation power by incorporating thousands of newly processed data from functional genomic assays in GRCh38 assembly, and now include probabilistic scores from the SURF algorithm that was the top performing non-coding variant predictor in CAGI 5^2^. We also provide interactive charts and genome browser views to allow users an easy way to perform exploratory analyses in different tissue contexts.

## Main Text

The update of RegulomeDB now includes > 650 million and > 1.5 billion genomic intervals in hg19 and GRCh38, respectively, which is in total five times the number in the previous version (Supplementary Fig. 2). Part of the data growth comes from over 3,000 new transcription factor (TF) ChIP-seq and chromatin accessibility experiments that were included from the ENCODE 3 and 4 phases ^3^. We also produced a comprehensive set of footprint predictions using a combination of over 800 chromatin accessibility experiments (642 from DNase-seq assays and 218 from ATAC-seq assays) and 591 TF motifs in GRCh38 as input to the TRACE pipeline^4^. In addition, we refined the included TF motifs by using the non-redundant vertebrates set from the JASPAR database^5^. We also integrated approximately 71 million variant-gene pairs in eQTL studies from the GTEx project^6^, and 450 thousand caQTLs (chromatin accessibility QTLs) from 9 recent publications (see Supplementary Note). Finally, we included chromatin state annotations from chromHMM in 833 biosamples^7^.

RegulomeDB accepts any query variants on the whole genome in either GRCh38 or hg19 genome assembly. The query variants can then be prioritized by functional prediction scores shown in a sortable table (Supplementary Fig. 3). For any variant of interest, an information page on five types of supported genomic evidence, as well as a genome browser view is displayed. Each of the six sections can be clicked to show more details on the genomic hits for functionality exploration (Supplementary Note; Supplementary Fig. 4 and 5).

RegulomeDB enables researchers to quickly separate functional variants from a large pool of variants by examining various assays and predictions, and assign tissue or organ specificity for each functional variant. Here we showcase this using six verified variants from recent literature^8–16^, and demonstrate the applicability of RegulomeDB to annotate those variants based on various sources of data (Fig. 1).

**Figure 1.**
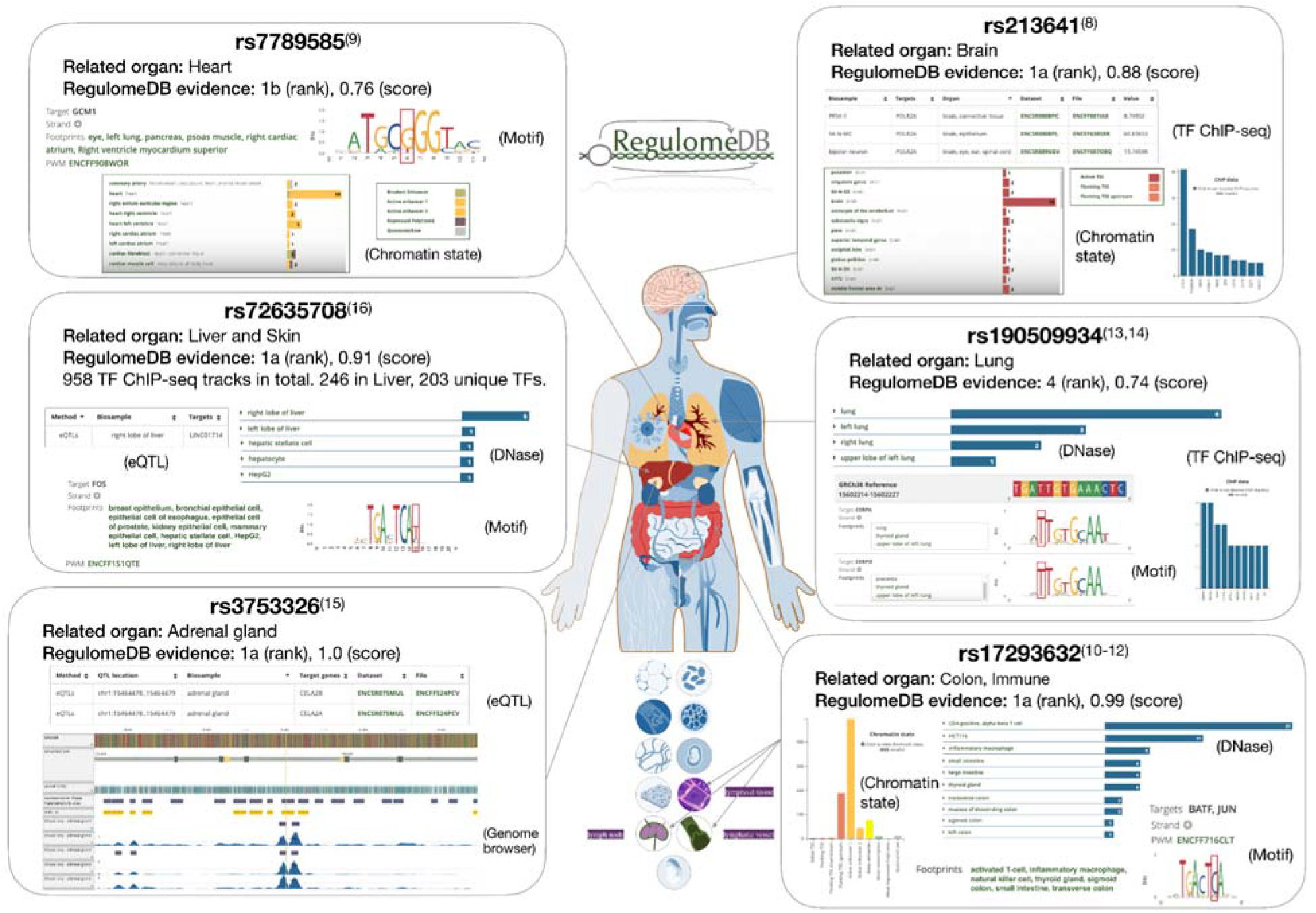
Prioritization of Functional Variants with RegulomeDB Version 2.1. Variants were identified by rsID from dbSNP in the title of each box. Supporting literature for characterizing their functionalities and related organs was superscripted in parentheses. Human body map was generated from ENCODE^3^.

TF motifs and ChIP-seq data together provide evidence about how a variant is likely to affect phenotype in a cell specific context. For example, rs213641 is known to affect behavioral responses to fear and anxiety stimuli^8^. The POLR2A binding and the active transcriptional start site (TSS) state in the brain indicate that rs213641 is likely to function in the brain through disrupting the TSS of STMN1. We also examined rs7789585 where RegulomeDB TF motif evidence suggests that mutation to the reference allele G would disrupt the binding of GCM1, which may interrupt the active enhancer state at the locus in the heart. Hocker and colleagues recently confirmed this hypothesis using reporter assays, and discovered that rs7789585 disrupts a KCNH2 enhancer and affects cardiomyocyte electrophysiologic function^9^. Another well-studied SNP, rs17293632, associated with Crohn’s disease and inflammatory bowel disease^10,11^ has strong AP-1 family motif evidence in RegulomeDB, implicating minor allele T in disease risk by reducing SMAD3 gene expression affecting the TGF-beta signaling pathway to regulate T cell activation and metabolism^12^.

DNase-seq assays and underlying footprint predictions identify open chromatin regions with mapped TF binding sites in hundreds of biosamples and can also be used to assign putative function to variants. rs190509934 has been associated with COVID-19 infection risk by affecting ACE2 expression level^13^. RegulomeDB shows hits to multiple DNase-seq peaks in lung related biosamples. Furthermore, RegulomeDB extends this tissue effect with the hypothesis that ACE2 expression level may be regulated by CEBP by its overlap with DNase footprints in the lung found in the upstream promoter region of ACE2^14^. Variants that are eQTLs provide correlation evidence for their target genes. rs3753326 is predicted as a regulatory variant affecting CELA2A gene expression level in the adrenal gland^15^.

RegulomeDB provides a user-friendly tool to annotate and prioritize variants in non-coding regions of the human genome, which can aid variant function interpretation and guide follow-up experiments. The latest updates of our pipeline facilitate the processing of BED formatted files and make it straightforward to integrate new datasets, such as peaks from ATAC-seq and 3D chromatin structure from Hi-C experiments. Meanwhile, the growing data from functional genomics assays enable the interpretation of variant functions in specific contexts. We plan to provide organ-specific prediction scores in a future version. We also welcome user feedback through the.

## Data availability

RegulomeDB v2.1 can be accessed through the web server at https://regulomedb.org. All datasets collected in RegulomeDB are accessible through the ENCODE portal: https://www.encodeproject.org/search/?internal_tags=RegulomeDB_2_1.

## Code availability

The code RegulomeDB uses is available on GitHub repository at https://github.org/ENCODE-DCC/regulome-encoded.

## References

1. Boyle, A. P. et al. Annotation of functional variation in personal genomes using RegulomeDB. Genome Res. 22, 1790–1797 (2012).

2. Dong, S. & Boyle, A. P. Predicting functional variants in enhancer and promoter elements using RegulomeDB. Hum. Mutat. 40, 1292–1298 (2019).

3. ENCODE Project Consortium et al. Expanded encyclopaedias of DNA elements in the human and mouse genomes. Nature 583, 699–710 (2020).

4. Ouyang, N. & Boyle, A. P. TRACE: transcription factor footprinting using chromatin accessibility data and DNA sequence. Genome Res. 30, 1040–1046 (2020).

5. Fornes, O. et al. JASPAR 2020: update of the open-access database of transcription factor binding profiles. Nucleic Acids Res. 48, D87–D92 (2020).

6. GTEx Consortium. The GTEx Consortium atlas of genetic regulatory effects across human tissues. Science 369, 1318–1330 (2020).

7. Boix, C. A., James, B. T., Park, Y. P., Meuleman, W. & Kellis, M. Regulatory genomic circuitry of human disease loci by integrative epigenomics. Nature 590, 300–307 (2021).

8. Brocke, B. et al. Stathmin, a gene regulating neural plasticity, affects fear and anxiety processing in humans. Am. J. Med. Genet. B Neuropsychiatr. Genet. 153B, 243–251 (2010).

9. Hocker, J. D. et al. Cardiac cell type–specific gene regulatory programs and disease risk association. Science Advances 7, eabf1444 (2021).

10. Devenish, L. P., Mhlanga, M. M. & Negishi, Y. Immune Regulation in Time and Space: The Role of Local- and Long-Range Genomic Interactions in Regulating Immune Responses. Front. Immunol. 12, 662565 (2021).

11. Miller, C. L. et al. Integrative functional genomics identifies regulatory mechanisms at coronary artery disease loci. Nat. Commun. 7, 12092 (2016).

12. Gate, R. E. et al. Genetic determinants of co-accessible chromatin regions in activated T cells across humans. Nat. Genet. 50, 1140–1150 (2018).

13. Horowitz, J. E. et al. Genome-wide analysis provides genetic evidence that ACE2 influences COVID-19 risk and yields risk scores associated with severe disease. Nat. Genet. 54, 382–392 (2022).

14. Beacon, T. H., Delcuve, G. P. & Davie, J. R. Epigenetic regulation of ACE2, the receptor of the SARS-CoV-2 virus1. Genome 64, 386–399 (2021).

15. Esteghamat, F. et al. CELA2A mutations predispose to early-onset atherosclerosis and metabolic syndrome and affect plasma insulin and platelet activation. Nat. Genet. 51, 1233–1243 (2019).

16. Kubota, N. & Suyama, M. An integrated analysis of public genomic data unveils a possible functional mechanism of psoriasis risk via a long-range ERRFI1 enhancer. BMC Med. Genomics 13, 8 (2020).

