## Supplemental Note for "Annotating and prioritizing human non-coding variants with RegulomeDB"

### Supplementary Note

### 1. Data sources

*Genomic variants*

The information of genomic variants was retrieved from dbSNP153 [1], including the positions and allele frequencies from different projects, such as the 1000 genome project [2], TOPMED [3] and GnomAD [4].

*ChIP-seq and DNase-seq*

We collected the peaks of ChIP-seq targeting transcription factors (TF) and DNase-seq experiments called by uniform pipeline from the latest release of the ENCODE project.

*PWM matching*

We downloaded the PWMs (position weight matrices) of 746 non-redundant TF motifs from JASPAR 2020 database [5].The kmers matching to TF motifs were called by TFM P-value with a threshold at $4^{-8}$ for each PWM [6]. Bowtie was used to map the kmers on the genome to determine the final PWM matching positions for the TF motifs [7]. The information content from each PWM was also integrated into the database and used as a feature to calculate the probabilistic score from the random forest model.

*Footprints*

Footprints were predicted with signals from 642 DNase-seq experiments and 591 TF motifs by the *TRACE* pipeline: https://www.encodeproject.org/search/?type=Annotation&internal_tags=RegulomeDB_2_1&annotation_type=footprints&software_used.software.name=trace[8]. TRACE is a computational method that incorporates DNase-seq signals and PWMs within a multivariate hidden Markov model to detect footprint regions with matching motifs.

*Chromatin states*

Chromatin states in 833 biosamples were called from chromHMM in EpiMap [9], and were directly retrieved from the ENCODE portal.

*eQTLs*
The eQTLs from the GTEx project across 49 human tissues were downloaded from the GTEx portal (https://storage.googleapis.com/gtex_analysis_v8/single_tissue_qtl_data/GTEx_Analysis_v8_eQTL.tar) [10]. The variant-gene pairs with the corresponding tissue were added as annotations in the database.

*caQTLs*
The chromatin accessibility QTLs (caQTLs) were collected from 9 publications [11–19] https://www.encodeproject.org/search/?type=Annotation&internal_tags=RegulomeDB_2_1&annotation_type=caQTLs. Only SNVs were included and lifted over from hg19 to GRCh38 if necessary [20].

*Prediction scores*
We provide a heuristic ranking and a probabilistic score for each query variant representing its potential of being a functional variant in regulatory elements. The heuristic ranking is defined in the same way as in the previous version of RegulomeDB [21]. The probabilistic score is calculated from a random forest model, TURF, trained with allele-specific TF binding SNVs [22]. We used a simplified version here only including binary features from functional genomic evidence as used in the heuristic ranking, as well as numeric features from information content in matched PWMs. We will include the whole feature set in a future release.

### 2. Database and web server design

RegulomeDB annotates a variant by intersecting its position with genomic intervals identified from a massive number of experiments and computational approaches. The database directly integrates the datasets from the ENCODE portal creating a genomic data service (https://github.org/ENCODE-DCC/genomic-data-service). The genomic intervals are parsed from BED formatted files and associated with metadata of the source experiments and computational pipelines from the ENCODE portal. These BED files are then indexed in Elasticsearch (https://www.elastic.co/) as in integer range type to enable efficient search against a query position. In total, over two billion genomic intervals representing ChIP-seq and DNase-seq peaks, matches to PWMs and DNase footprints, eQTLS, caQTLs and chromatin states are indexed in Elasticsearch. After each search, the JSON objects associated with the intersected intervals are returned and passed on to generate ranking scores from RegulomeDB 1.1 and new probabilistic scores from TURF [22,23]. The query results are displayed with a web interface (https://github.org/ENCODE-DCC/regulome-encoded) that contains charts and interaction figures, which can be customized by users.

### 3. New interface for variant functionality exploration

The RegulomeDB v2.1 web server accepts any query variant on the whole genome in either GRCh38 or hg19 genome assembly. A toggle above the search box allows users to switch between the two assemblies. The search box allows any user to input multiple queries (up to 500 at a time) (Supplementary Figure 3). The input query can be in three formats: 1) rsID (from dbSNP database v153); 2) chromosome position for a single nucleotide variant; 3) chromosome position for a chromosome region. In the third case, all variants on the chromosome region at >1% allele frequency from dbSNP153 will be queried. The backend then intersects the variant(s) position with the genomic intervals of annotations obtained predicted from functional genomics experiments and returns a sortable summary table of variant scores (Supplementary Figure 3), including a ranking score and a probabilistic score showing its potential of being a regulatory variant. In addition, a dbSNP rsID will link to the query variant if it exists.

After clicking on any field of a row in the score table, a more detailed information page on genomic evidence is shown for the variant of interest (Supplementary Figure 4, Supplementary Figure 5). The top of the page shows some basic information on the variant position, scores, and allele frequencies from the dbSNP database. While on the bottom is the initial summary section on genomic annotations' hits. Since a single query can hit up to 2,000 results, the initial summary section is divided into five data types; TF binding sites from ChIP-seq, chromatin states from chromHMM, chromatin accessibility, PWM matching or footprint predictions, and eQTLs or caQTLs. In addition, a genome browser section is also available to view the specific DNase-seq and ChIP-seq data, which can aid in variant interpretation.

Each of the six sections can be clicked to display more details on the genomic hits from specific assays, such as the biosample of DNase peaks and the transcription factors of ChIP-seq peaks. The chromatin state tab shows the chromHMM state for each of the 833 biosamples, which also includes an intuitive body map colored by the most active chromatin state in each organ. Furthermore, the genome browser tab provides an interaction view for exploring the gene transcripts along with DNase-seq and ChIP-seq peaks near the variant of interest (shown as a yellow highlight). The tracks on the genome browser can be further filtered using a modal that allows one to sub-select by specific organ/cell types, biosample types, file types, assay methods, or by TF targets.

**References**

1. Sherry, S. T. dbSNP: the NCBI database of genetic variation. *Nucleic Acids Research* vol. 29 308–311 Preprint at https://doi.org/10.1093/nar/29.1.308 (2001).

2. 1000 Genomes Project Consortium *et al.* A global reference for human genetic variation. *Nature* **526**, 68–74 (2015).

3. Taliun, D. *et al.* Sequencing of 53,831 diverse genomes from the NHLBI TOPMed Program. *Nature* **590**, 290–299 (2021).

4. Karczewski, K. J. *et al.* The mutational constraint spectrum quantified from variation in

141,456 humans. *Nature* **581**, 434–443 (2020).

5. Fornes, O. *et al.* JASPAR 2020: update of the open-access database of transcription factor binding profiles. *Nucleic Acids Res.* **48**, D87–D92 (2020).
6. Touzet, H. & Varré, J.-S. Efficient and accurate P-value computation for Position Weight Matrices. *Algorithms Mol. Biol.* **2**, 15 (2007).
7. Langmead, B., Trapnell, C., Pop, M. & Salzberg, S. L. Ultrafast and memory-efficient alignment of short DNA sequences to the human genome. *Genome Biol.* **10**, R25 (2009).
8. Ouyang, N. & Boyle, A. P. TRACE: transcription factor footprinting using chromatin accessibility data and DNA sequence. *Genome Res.* **30**, 1040–1046 (2020).
9. Boix, C. A., James, B. T., Park, Y. P., Meuleman, W. & Kellis, M. Regulatory genomic circuitry of human disease loci by integrative epigenomics. *Nature* **590**, 300–307 (2021).
10. GTEx Consortium. The GTEx Consortium atlas of genetic regulatory effects across human tissues. *Science* **369**, 1318–1330 (2020).
11. Degner, J. F. *et al.* DNase I sensitivity QTLs are a major determinant of human expression variation. *Nature* **482**, 390–394 (2012).
12. Schwartzentruber, J. *et al.* Molecular and functional variation in iPSC-derived sensory neurons. *Nat. Genet.* **50**, 54–61 (2018).
13. Khetan, S. *et al.* Type 2 Diabetes–Associated Genetic Variants Regulate Chromatin Accessibility in Human Islets. *Diabetes* **67**, 2466–2477 (2018).
14. Gate, R. E. *et al.* Genetic determinants of co-accessible chromatin regions in activated T cells across humans. *Nat. Genet.* **50**, 1140–1150 (2018).
15. Tehranchi, A. *et al.* Fine-mapping cis-regulatory variants in diverse human populations. *Elife* **8**, (2019).
16. Kumasaka, N., Knights, A. J. & Gaffney, D. J. High-resolution genetic mapping of putative causal interactions between regions of open chromatin. *Nat. Genet.* **51**, 128–137 (2019).
17. Zhao, Q. *et al.* Molecular mechanisms of coronary disease revealed using quantitative trait

loci for TCF21 binding, chromatin accessibility, and chromosomal looping. *Genome Biol.* **21**, 135 (2020).

18. Liang, D. *et al.* Cell-type-specific effects of genetic variation on chromatin accessibility during human neuronal differentiation. *Nat. Neurosci.* **24**, 941–953 (2021).
19. Currin, K. W. *et al.* Genetic effects on liver chromatin accessibility identify disease regulatory variants. *Am. J. Hum. Genet.* **108**, 1169–1189 (2021).
20. Kuhn, R. M., Haussler, D. & Kent, W. J. The UCSC genome browser and associated tools. *Brief. Bioinform.* **14**, 144–161 (2013).
21. Boyle, A. P. *et al.* Annotation of functional variation in personal genomes using RegulomeDB. *Genome Res.* **22**, 1790–1797 (2012).
22. Dong, S. & Boyle, A. P. Prioritization of regulatory variants with tissue-specific function in the non-coding regions of human genome. *Nucleic Acids Res.* (2021) doi:10.1093/nar/gkab924.
23. Dong, S. & Boyle, A. P. Predicting functional variants in enhancer and promoter elements using RegulomeDB. *Hum. Mutat.* **40**, 1292–1298 (2019).

### Supplementary Figures

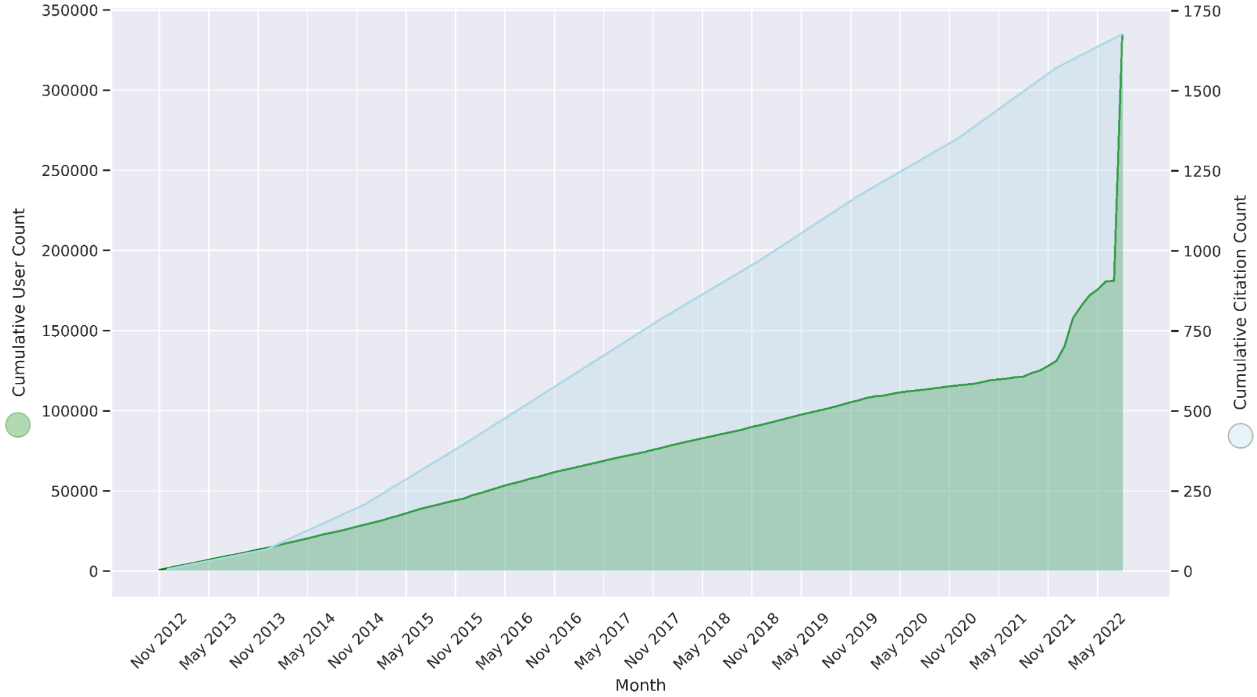


**Supplementary Figure 1.** Popularity of RegulomeDB.

The x-axis is month and year since RegulomeDB first published in 2012. The left y-axis is cumulative user count (green). The right y-axis is cumulative citation count (light blue). The citation count data are derived from Clarivate Web of Science. © Copyright Clarivate 2022. All rights reserved.

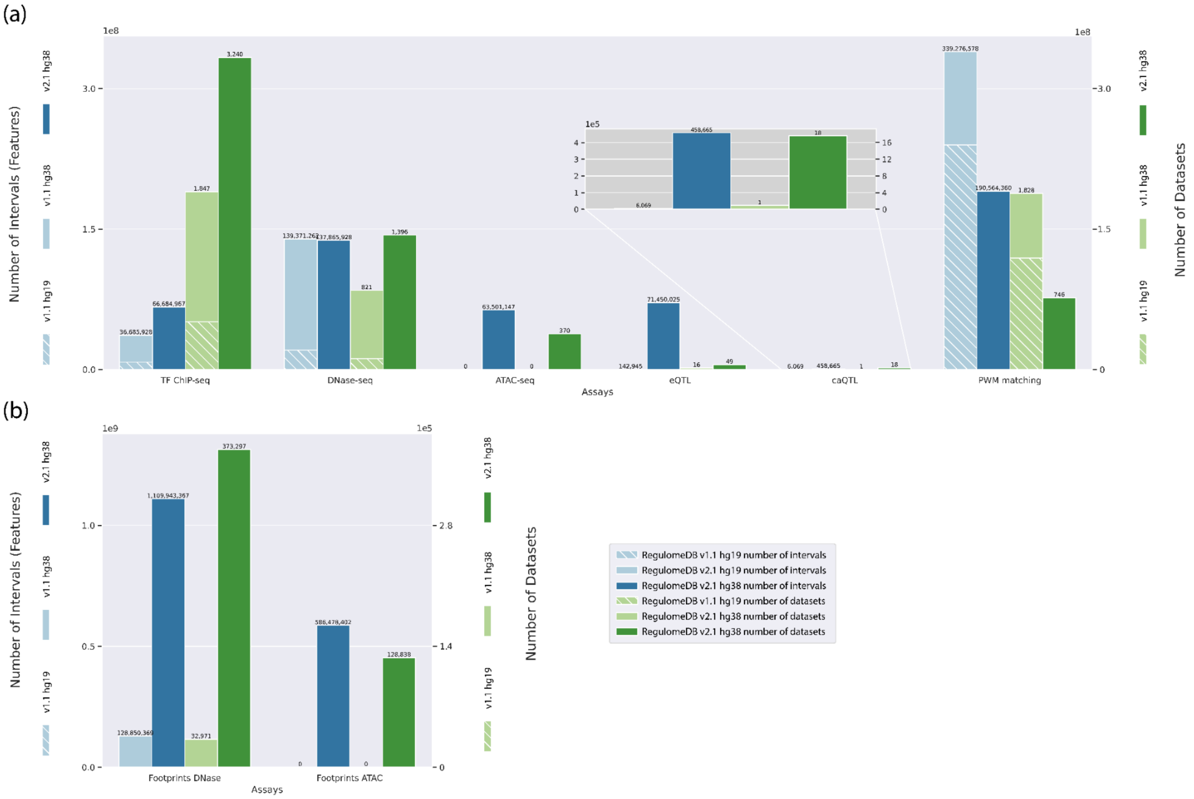


**Supplementary Figure 2.** Overview of RegulomeDB Version 2.1 Data Growth and Refinement. Statistics on database content. Numbers under each data type include all experiments across different treatment conditions and biosamples. All numbers are RegulomeDB v2.1 stats, in hg19 or hg38.

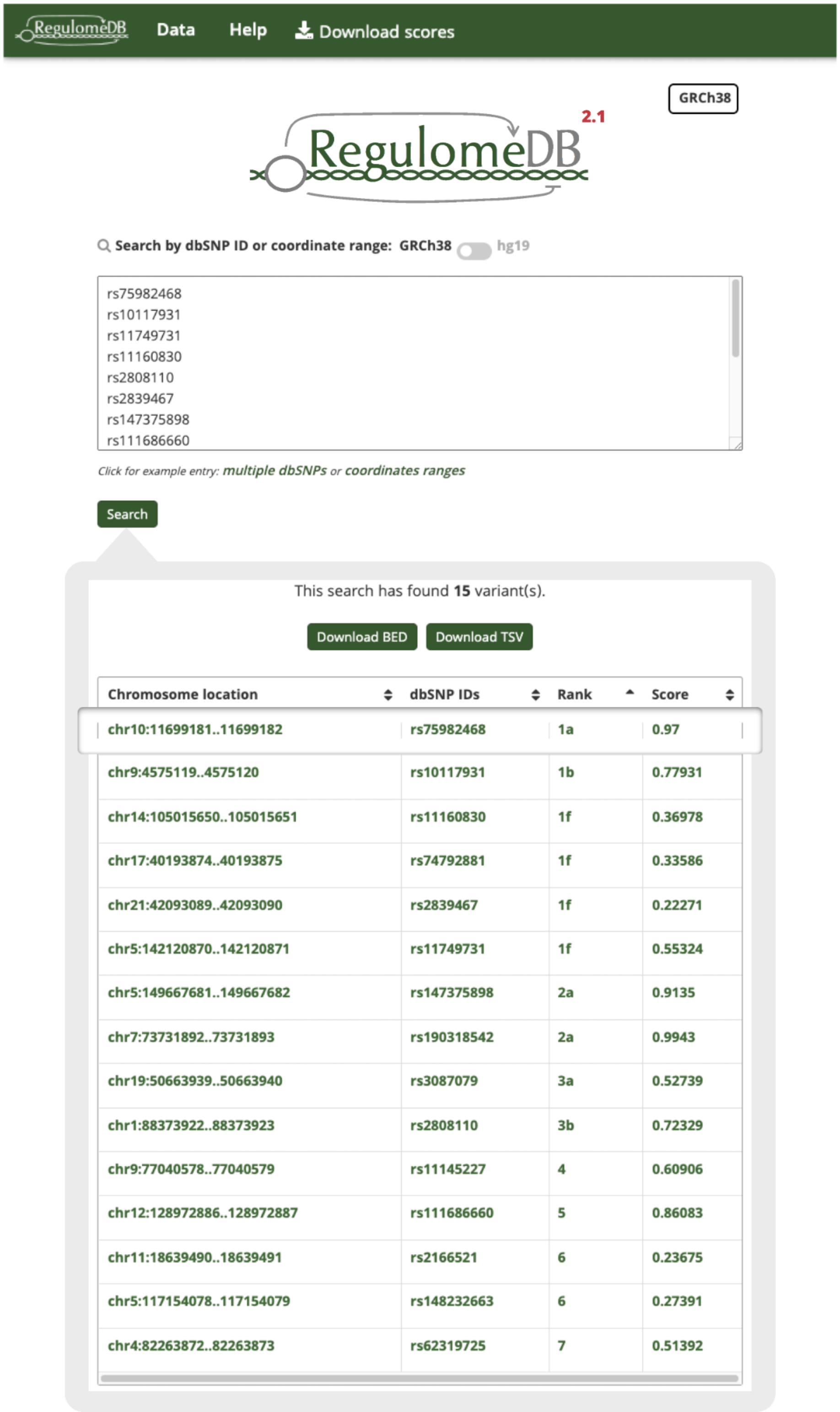


| Chromosome location | dbSNP IDs | Rank | Score |
|---|---|---|---|
| chr10:11699181..11699182 | rs75982468 | 1a | 0.97 |
| chr9:4575119..4575120 | rs10117931 | 1b | 0.77931 |
| chr14:105015650..105015651 | rs11160830 | 1f | 0.36978 |
| chr17:40193874..40193875 | rs74792881 | 1f | 0.33586 |
| chr21:42093089..42093090 | rs2839467 | 1f | 0.22271 |
| chr5:142120870..142120871 | rs11749731 | 1f | 0.55324 |
| chr5:149667681..149667682 | rs147375898 | 2a | 0.9135 |
| chr7:73731892..73731893 | rs190318542 | 2a | 0.9943 |
| chr19:50663939..50663940 | rs3087079 | 3a | 0.52739 |
| chr1:88373922..88373923 | rs2808110 | 3b | 0.72329 |
| chr9:77040578..77040579 | rs11145227 | 4 | 0.60906 |
| chr12:128972886..128972887 | rs111686660 | 5 | 0.86083 |
| chr11:18639490..18639491 | rs2166521 | 6 | 0.23675 |
| chr5:117154078..117154079 | rs148232663 | 6 | 0.27391 |
| chr4:82263872..82263873 | rs62319725 | 7 | 0.51392 |

**Supplementary Figure 3.** RegulomeDB Query Interface.
An example query with the rsIDs of variants from dbSNP database. Upon clicking the search buttons, a summary table representing prediction scores for all query variants will be displayed. (See Supplementary Note 3 for more details.)

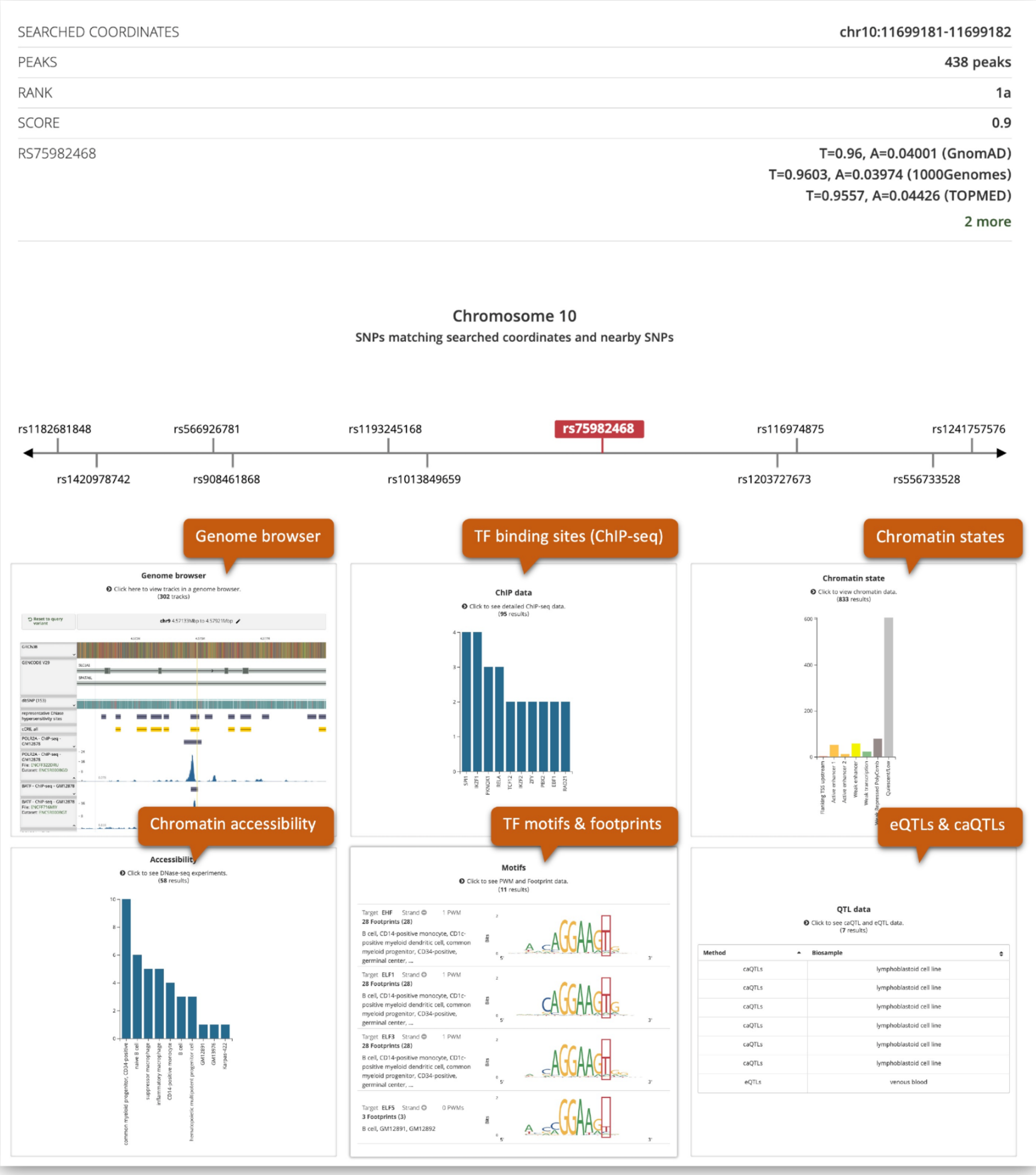


**Supplementary Figure 4.** RegulomeDB Result Overview Page of rs75982468.

For any variant of interest, a results page with more information on the hits of genomic annotations is available. Each of the six sections at the bottom can be clicked to expand more details on each data type.

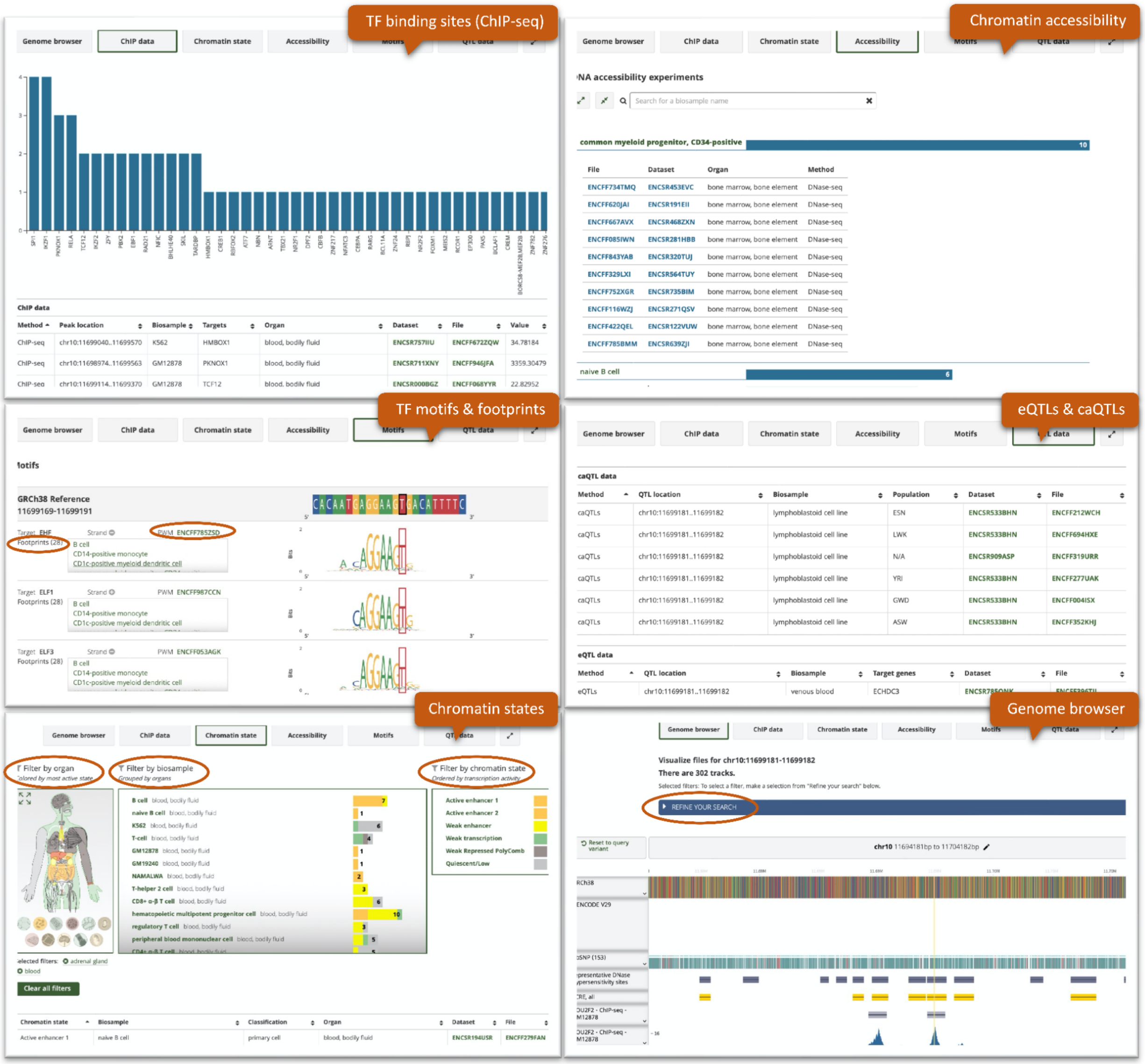


**Supplementary Figure 5.** RegulomeDB Expanded Pages of rs75982468.

The expanded pages of each section shows details on the genomic experiments and annotations, such as the biosample, organ, TF target and the peak file called from the ENCODE project. The body map under the chromatin states view is colored by the most active state among all biosamples in each organ, which gives an intuitive way to explore the candidate organs where the query variant might be functional. Users can also explore the nearby genes of the query variant under the genome browser view.
